# Food-Allergic Children Harbour a Reduced Oral Bacterial Load, Giving Misleading 16S rRNA Amplicon Profiles

**DOI:** 10.64898/2026.02.10.705039

**Authors:** Simona Panelli, Lodovico Sterzi, Clara Bonaiti, Miriam Acunzo, Enza Carmina D’Auria, Gianvincenzo Zuccotti, Francesco Comandatore

## Abstract

**Background:** Food allergies have become a major public health concern in Western countries due to their rising prevalence. The microbiota is increasingly recognized as a key factor in their pathogenesis. Using V3–V4 16S rRNA amplicon sequencing, our group previously reported alterations in the oral microbiota of children with food allergies compared to controls. Given growing awareness of biases associated with relative-abundance analyses in 16S rRNA studies, we re-analyzed the salivary DNA samples from our earlier work.

**Methods:** Total bacterial loads were quantified by qPCR using universal 16S rRNA primers, enabling the estimation of absolute abundances for each taxon in each sample.These data were then used to reassess key microbial community features and to directly compare conclusions derived from absolute versus relative abundance approaches.

**Results:** Allergic children exhibited a marked reduction in salivary bacterial load, a feature not detected in the original relative-abundance analysis. Incorporating absolute abundances also partially reshaped the previously defined taxonomic pictures. Similar distortions affected functional predictions, such as Short Chain Fatty Acids (SCFA)-producing capacity.

**Conclusions:** Our data confirm that variations of the bacterial load across samples hinder efforts to associate microbiota features and disease phenotypes. We identify reduced oral bacterial load as a previously unrecognized characteristic of children with food allergies. This observation aligns with reports in other immune-mediated disorders, suggesting that overall microbiota depletion may represent a defining and potentially diagnostically feature of disease-associated ecosystems.

## INTRODUCTION

Food allergies (FA) are inappropriate immune reactions triggered by the ingestion of normally innocuous food antigens (allergens), resulting in local or systemic reactions. Symptoms range from life threatening conditions, such as anaphylaxis, to respiratory, cutaneous and gastrointestinal disorders^[1]^. FA prevalence can reach up to 10%, with considerable variation across geographical regions, allergens and age groups^[2]^. Over the past century, FA prevalence has significantly increased in western countries^[3]^ emerging as a major public health concern.

The growing prevalence supports the involvement of environmental factors in the pathogenesis of FA. Indeed, like the other allergic diseases, FA are complex conditions resulting from the interplay of multiple endogenous and exogenous influences that can either attenuate or exacerbate their pathogenesis^[4,5]^. Although an exact genetic basis has yet to be fully elucidated, recent evidence highlights the involvement of the epigenome as a key mediator linking environmental exposures to disease susceptibility. Early-life factors, pollution, chemicals, diet, infections and interactions with other organisms have all been implicated in the pathogenesis^[6]^. Microbial exposure, in particular, is now recognized as a critical environmental risk modifier, with both pathogens and commensal bacteria generally endowed with protective roles^[7,8]^. Recent works point to a central role for the microbiota^[8]^, with its influence manifesting through multiple mechanisms. For example the microbiota exerts immunomodulatory effects *via* the production of metabolites such as short chain fatty acids (SCFA), which promote tolerance, possess anti-inflammatory activities and contribute to the epigenetic regulation of the immune system^[9]^. A dysregulated homeostatic interaction between the host and its microbiota has been identified as a predisposing factor for FA, with microbial dysbiosis preceding the onset of food sensitization and playing a critical role in the establishment, or failure, of oral tolerance ^[8]^.

Recent studies have highlighted the pivotal role of the oral host-commensal environment ^[10]^. Using amplicon sequencing on the V3-V4 regions of the 16S rRNA gene, our group reported alterations in the oral bacterial taxonomic profile of children with food allergies compared to healthy controls ^[11]^. In agreement with previous data ^[10]^, several taxa resulted significantly depleted in the allergic group, including the genus *Streptococcus* and its higher taxonomic rankings up to the phylum *Firmicutes*; the genus *Neisseria* and the corresponding family (*Neisseriaceae*), order (*Neisseriales*) and class (*Betaproteobacteria*); as well as the genera *Prevotella* and *Veillonella*. However, our results revealed that the most statistically significant difference between the two groups was a marked enrichment, in allergic children, of sequences belonging to “*Candidatus* Gracilibacteria”. This lineage belongs to the Candidate Phyla Radiation (CPR), also known as *Patescibacteria*, a peculiar and still poorly characterized bacterial clade ^[12]^. In addition, a second *Patescibacteria* lineage, “*Candidatus* Saccharimonadia” (formerly TM7 or “*Candidatus* Saccharibacteria”), exhibited a trend to increase in the allergic group. Members of *Patescibacteria*, “*Ca*. Saccharimonadia” in particular, are increasingly associated with inflammatory diseases, and growing evidence points to their involvement in immunomodulatory processes ^[13]^. Although their relevance to host physiology and disease is becoming progressively recognized, these organisms remain largely understudied for several methodological and biological reasons ^[12]^. Together, these considerations prompted us to further investigate the enrichment of *“Ca. Saccharimonadia”* and other CPR lineages in order to better contextualize their role within the oral microbiota of children with food allergy (FA).

Another key premise for refining our results came from a recent work by Bruijning and colleagues (2023) ^[14]^ which highlighted a potential source of bias in the way microbiota data are typically presented in 16S rRNA amplicon sequencing studies. Indeed, 16S rRNA amplicon sequencing can not provide the absolute abundance of bacterial taxa in a sample, but it expresses it as a percentage. The authors evidenced that the using of relative abundances can bias the comparison of taxa abundance and of their changing, because percentages are inherently autocorrelated ^[15]^. In other words, because the sum of relative abundances in a sample must equal one, a decrease in the relative abundance of one taxon necessarily results in an apparent increase in another, regardless of their actual quantities. This can introduce distortions, particularly when assessing correlation between variables, or identifying taxonomic signatures across treatment groups ^[16,17]^. Bruijning and colleagues (2023) ^[14]^ propose the use of absolute abundances, rather than relative, as a way to alleviate these distortions. One way they propose is integrating microbial relative abundance data with estimates of the total bacterial load of the sample, derived from qPCR estimates.

Considering that studies comparing absolute vs. relative abundances are beginning to emerge ^[18,19]^, we decided to re-analyze salivary DNA samples we previously characterized by 16S V3-V4 amplicon sequencing in our earlier food allergy study ^[11]^. For each sample, we estimated the total bacterial load *via* qPCR with universal bacterial primers targeting the 16S rRNA gene. This enabled us to re-analyze the microbiota data, estimating the absolute abundances of each taxon in each sample. The new results only partially overlap those based on relative abundances and, notably, they reveal a variation in total bacterial counts between the allergic and control groups that was not apparent from the relative frequency data. Our novel results open to a different view about the host-oral microbiota relationship in food allergy.

## METHODS

### Ethics statement

This study analyzed the collection of human salivary DNA samples (n=59) already characterized in a previous microbiota study on food allergy ^[11]^. The research was conducted according to the declaration of Helsinki, and all methods followed the relevant guidelines and regulations. The Ethics Committee of ASST-Fatebenefratelli-Sacco approved the study (Ref. n. 2021/ST/041). Privacy rights of subjects were carefully observed and authors did not have access to information that could identify individual participants. During the recruitment of subjects, parents signed a written informed consent to allow their child to participate in the study. Samples and data were first accessed for research purposes on 01 February 2022.

### Quantitative PCR (qPCR) for estimating total bacterial load

We used a pan-bacterial qPCR protocol on the 16S rRNA gene to estimate the total bacterial load of the 59 DNA samples previously characterized by D’Auria et al. ^[11]^. In that study, DNA was extracted from saliva of children suffering from food allergies (n=29) and controls (n=30), and subjected to 16S rRNA V3-V4 amplicon sequencing. The characteristics of the cohort have been described in our previous work. For the present study, the same DNA tubes were used: samples were not re-extracted in order to avoid any kind of variation that would have distorted the comparison between the qPCR results and the amplicon sequencing analysis

As suggested by Bruijning and colleagues (2023) ^[14]^, we performed the total bacterial quantification *via* qPCR using pan-bacterial primers targeting the 16S rRNA gene. We selected the 926F/1062R ^[20]^ primer pair based on literature and on our previous *in silico* results, in which we assessed the general absence of bias in binding 16S rRNA genes from different portions of the bacterial tree, comprising the *Patescibacteria* ^*[21]*^.

Specifically, the number of 16S rRNA gene copies were determined in 2-μL DNA samples, in a total qPCR volume of 15μL, also containing 7.5 μL of 2x SsoAdvanced Universal SYBR® Green Supermix (BioRad, Hercules, CA), 4.7 μL of PCR amplification-grade water (Promega Corporation, Wisconsin, USA), and 0.4 μL of 10 μM stocks for the pan-bacterial primers 926F and 1062R ^[20]^. Each sample was qPCR-amplified in three technical replicates. The qPCR assays were performed on a BioRad CFX Connect real-time PCR System (BioRad, Hercules, CA) with the following profile: initial denaturation (95°C 5 min), 40 cycles at 95°C 15s, 61.5°C 15s, 72°C 20s. Finally, a melting curve was generated, with a range of temperature between 60° and 95°C.

### Data analysis

The total bacterial loads obtained by qPCR for allergic and control children were compared using the Wilcoxon–Mann–Whitney test, with a significance threshold (p-value) set to 0.05.

The absolute abundance of each taxon in each sample was then estimated by multiplying the total bacterial load (obtained by qPCR, as described above) by the relative frequencies obtained from the 16S V3-V4 amplicon sequencing retrieved from D’Auria et al. 2023^[11]^.

Taxonomic pictures and ecological indices of β-diversity obtained using both absolute and relative abundances were computed using the R library Vegan (https://CRAN.R-project.org/package=vegan). The α-diversity indices were not recalculated as they are not expected to change when considering absolute abundances, obtained by multiplying relative frequencies by a constant. Concerning the microbial profiles obtained by relative and absolute abundances, they were compared using Mann–Whitney U test, with a significance threshold (p-value) set to 0.05.

OTU sequences classified as Unclassified at phylum level were retrieved and BLASTn searched against the SILVA v138 database ^[22]^ to assess the closest phylum.

The production of the SCFAs for each sample (i.e. allergic and control) was finally investigated combining Picrust2 ^[23]^ predictions and, respectively, absolute and relative OTU abundances. The inferred SCFA production of allergic vs control children was then compared using Wilcoxon–Mann–Whitney test, with a significance threshold (p-value) set to 0.05.

## RESULTS

### qPCR estimation of total bacterial load in salivary DNA from allergic and control individuals

Using a pan-bacterial 16S rRNA qPCR-based approach, we estimated the total bacterial load in the 59 salivary DNA samples, collected from allergic and control children, previously characterized by D’Auria et al. ^[11]^. As shown in [Figure 1], allergic individuals displayed a significantly lower total bacterial load than controls (Wilcoxon–Mann–Whitney test, p-value < 2.2x10^-16^). In particular, the median bacterial quantification for 2-μl samples was 2.4x10^7^ for allergic children and 3.5x10^7^ for the control group, corresponding respectively to 1.2x10^9^/ml and 1.8x10^9^/ml.

**Figure 1.**
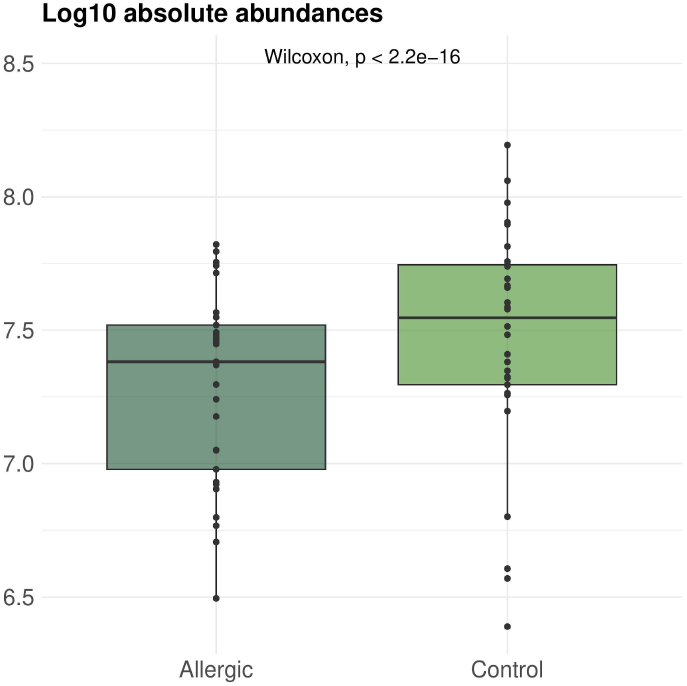
Absolute bacterial abundances in children suffering from food allergy and controls. Log10 of absolute bacterial abundance estimated as the 16S rRNA gene copies per 2-μl samples determined by pan-bacterial qPCR. Data for the two groups were compared using the Wilcoxon–Mann–Whitney test.

### Comparison between the ecological indexes of β-diversity computed from relative and absolute bacterial taxa abundances in allergic and control individuals

The Principal Coordinates Analysis (PCoA), computed both on relative and absolute abundances at the families and genera taxonomic levels, is reported in [Figure 2]. For all the four PCoAs, the PERMANOVA test coherently reported the statistically significant separation between the oral microbiota of allergic and control children.

**Figure 2.**
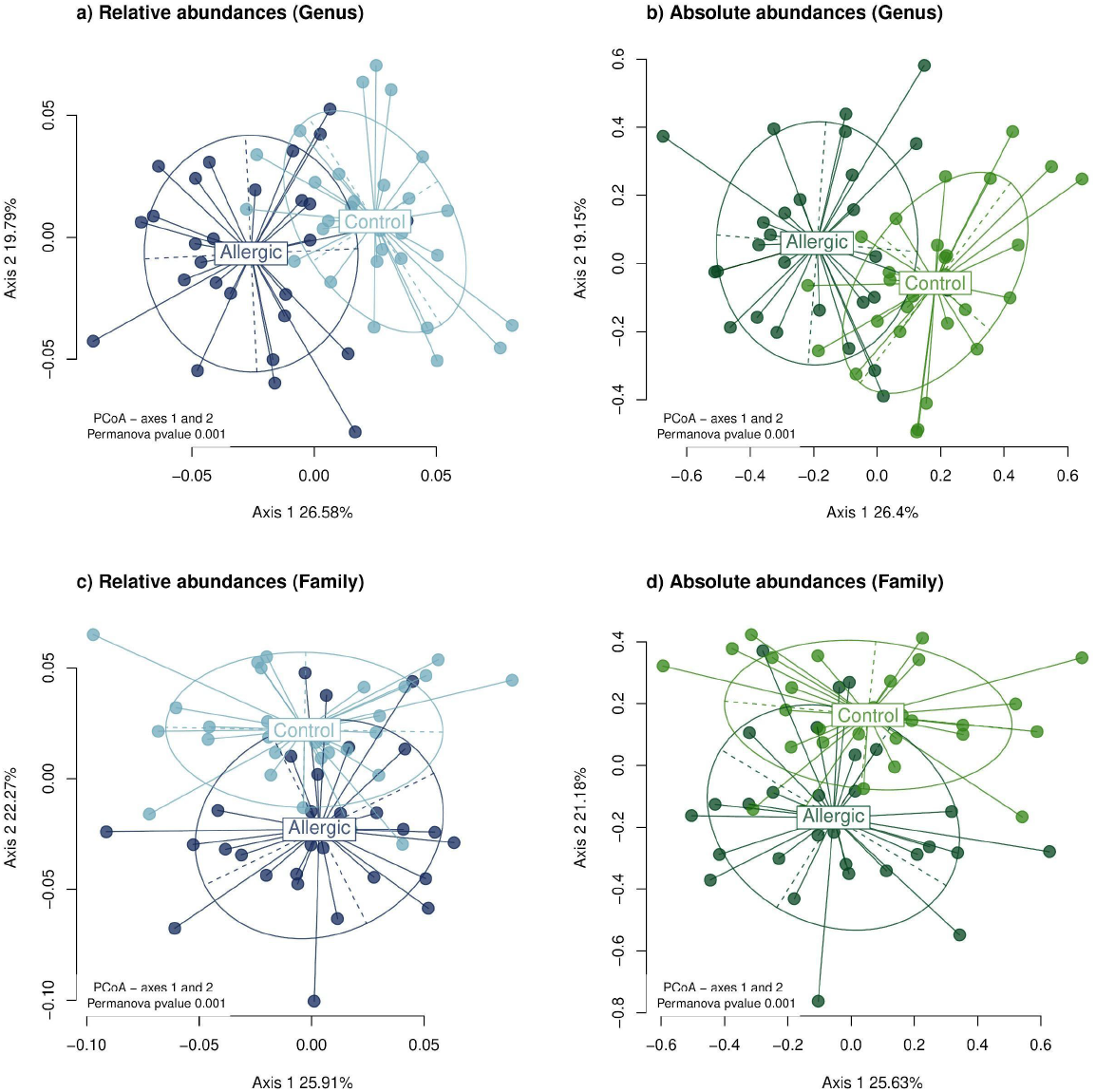
β-diversity analysis. The microbiota composition of allergic vs control individuals were compared by Principal Coordinates Analysis (PCoA). (a-b) PCoA at the taxonomic levels of genus; (c-d) PCoA at the taxonomic levels of family. PCoA obtained for the relative abundances are in plots a and c (samples from allergic children are colored in dark blue and those from controls in light blue) while those computed from the absolute abundances are shown in plots b and d (allergic in dark green and controls in light green). Allergic vs control microbiota compositions were compared using the PERMANOVA test and significant threshold was set < 0.05.

### Comparison between relative and absolute bacterial taxa abundances in allergic and control individuals

For each of the 59 saliva samples, the absolute abundance of each bacterial taxon was estimated combining the total bacterial load and the bacterial taxa relative abundances retrieved from D’Auria and colleagues (2023) ^[11]^ (see Material and Methods for details).

The 16S rRNA amplicon sequencing analysis identified a total of 303 bacterial taxa across all taxonomic levels. Among these, 217 taxa (72%) showed consistent patterns using relative and absolute abundances (Supplementary Table 1). For example, the phylum *Firmicutes* was significantly less abundant in allergic children than in controls, according to both relative and absolute abundances. Overall, coherent results for relative and absolute abundances were observed for six of 11 phyla (55%), 13/20 classes (65%), 33/47 orders (70%), 53/70 families (68%) and 112/147 genera (76.%) (Supplementary Table 1).

Among the 303 taxa identified, 156 showed significant differences in relative and/or absolute abundance between allergic and control children. These comprised six phyla, 10 classes, 24 orders, 45 families, and 71 genera. Of these, coherent differences between the two groups detected by both absolute and relative measures, were observed for one of the six phyla (17%), seven of the 10 classes (70%), 10 of the 24 orders (42%), 20 of the 45 families (44%), and 36 of the 71 genera (51%) (Supplementary Table 1).

Considering the taxonomic level of phyla, among the six mentioned above, *Firmicutes* was the only group showing concordant results, with lower relative and absolute abundances in allergic children compared to controls [Figure 3]. *Actinobacteriota, Bacteroidota*, and *Spirochetota* were reduced in allergic children only when considering absolute abundances. In contrast, *Patescibacteria* and unclassified Bacteria displayed higher relative abundances in allergic children, while their absolute abundances remained unchanged.

**Figure 3.**
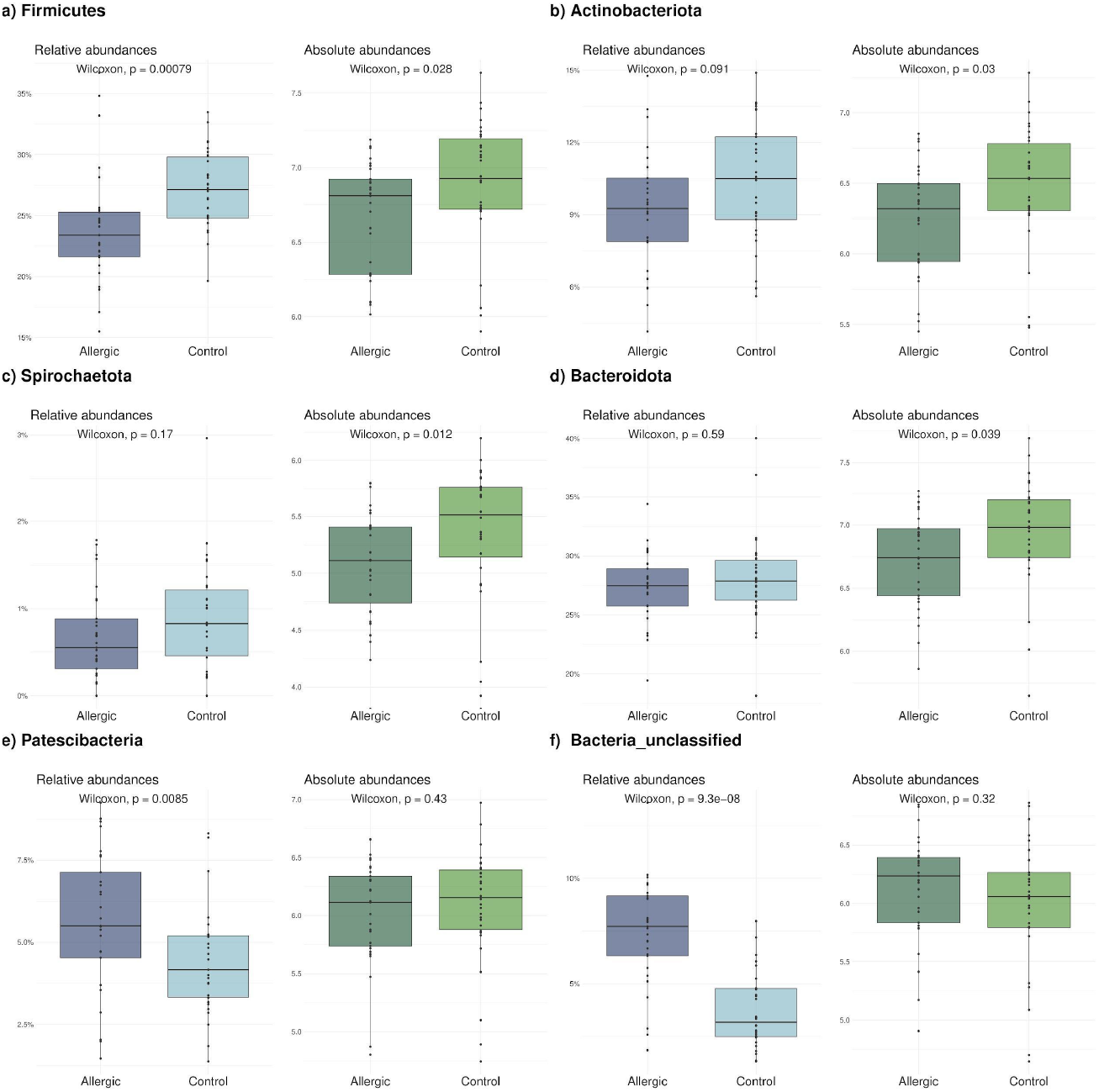
Plots a-f show relative and absolute abundances of bacterial phyla with significant differences in relative and/or absolute abundances between control and allergic children. In each plot, two couples of boxplots are shown: on the left, boxplots comparing the relative abundances between allergic (dark blue) and control (light blue) subjects; on the right, the boxplots comparing the absolute abundances between allergic (dark green) and control (light green) subjects.

*Firmicutes* was the only phylum that resulted significantly different in allergic children vs controls according to both absolute and relative abundances. As reported in Table S1, a total of 98 *Firmicutes* taxa were identified. Of these, 48 exhibited no significant differences between allergic and control children in either relative and absolute abundances, while 19 taxa yielded discordant results. The remaining 31 *Firmicutes* taxa exhibited significant between-group differences in both analyses. Of these 31 taxa, 27 were significantly reduced in allergic children, mirroring the overall phylum-level trend. These included, for example, the orders *Lactobacillales, Staphylococcales*, and *Lachnospirales*; the families *Aerococcaceae, Carnobacteriaceae, Gemellaceae*, and *Lachnospiraceae*; and the genera *Solobacterium, Granulicatella, Streptococcus*, and *Gemella*. The remaining four taxa were low-abundance, unclassified *Firmicutes* lineages, and were the only ones resulting significantly enriched in allergic children according to both relative and absolute abundances.

*Patescibacteria* and sequences unresolved by the 16S rRNA V3-V4 amplicon sequencing represented the portion of the oral microbiota stable in terms of absolute abundance in allergic children, despite the marked, overall reduction of the oral bacterial load in this group. The taxonomic identity of these unresolved sequences was explored as described in Materials and Methods. [Figure 5] shows the distribution of the phyla corresponding to their closest BLAST matches. As evident from the Figure, these sequences represent a heterogeneous set of bacterial lineages spanning a broad range of the known bacterial diversity. The most represented phylum is *Firmicutes*, for both allergic and control children, but several other phyla are also detected.

General patterns of concordance/discordance between absolute and relative abundances are shown in the heatmap of [Figure 4] for all taxonomic levels.

**Figure 4.**
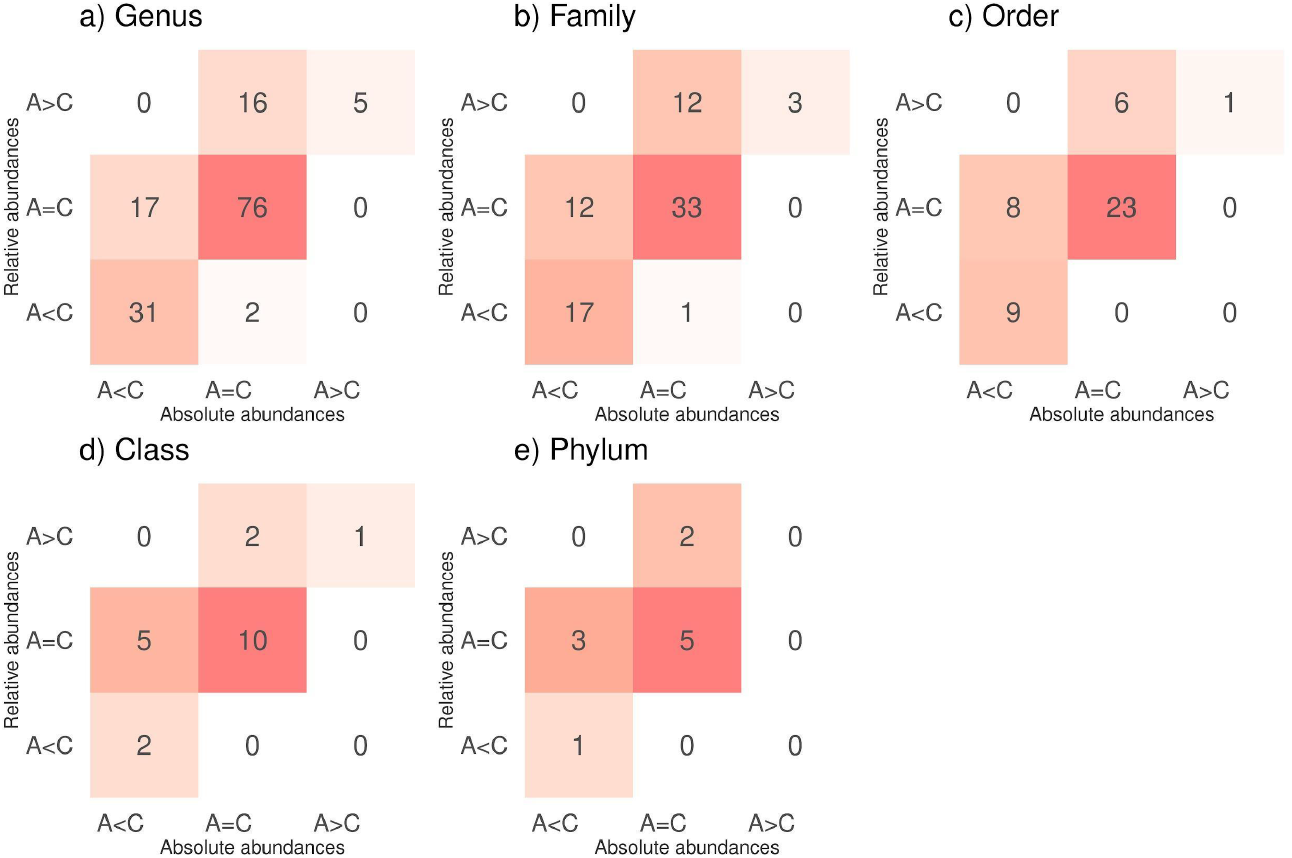
Heatmap showing patterns of concordance and discordance between relative and absolute abundances across all taxonomic levels. Rows represent relative abundance comparisons, and columns represent absolute abundance comparisons. Row and column labels refer to comparisons between allergic (A) and control (C) individuals: A > C indicates taxa with statistically significantly higher abundance in allergic individuals; A = C indicates no statistically significant difference; A < C indicates taxa with significantly lower abundance in allergic individuals compared with control individuals. Values within cells indicate the number of taxa exhibiting the pattern defined by each row–column combination.

**Figure 5.**
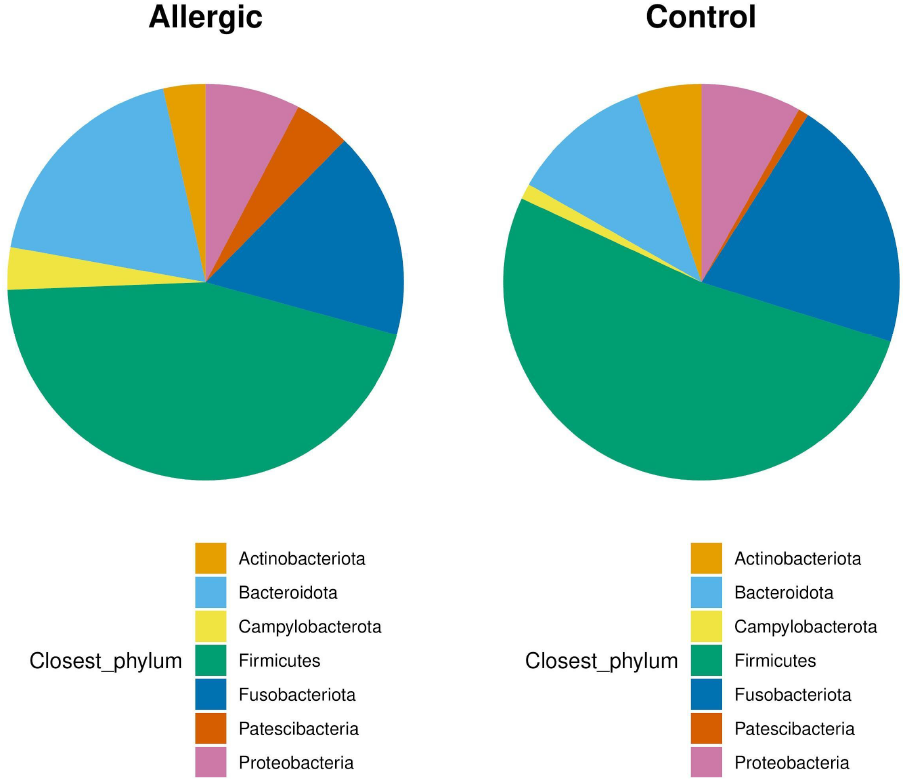
Distribution of the bacterial phyla closest to sequences unclassified at the phylum level following 16S rRNA V3-V4 amplicon sequencing. The allergic group is shown on the left, and the control group on the right.

### Prediction of SCFA production from relative and absolute bacterial taxa abundances in allergic and control individuals

The potential production of butyrate, propionate, and succinate from the oral microbiota of the two groups was inferred by integrating PICRUSt2 functional predictions with the absolute or relative abundances of the corresponding taxa [Figure 6].

**Figure 6.**
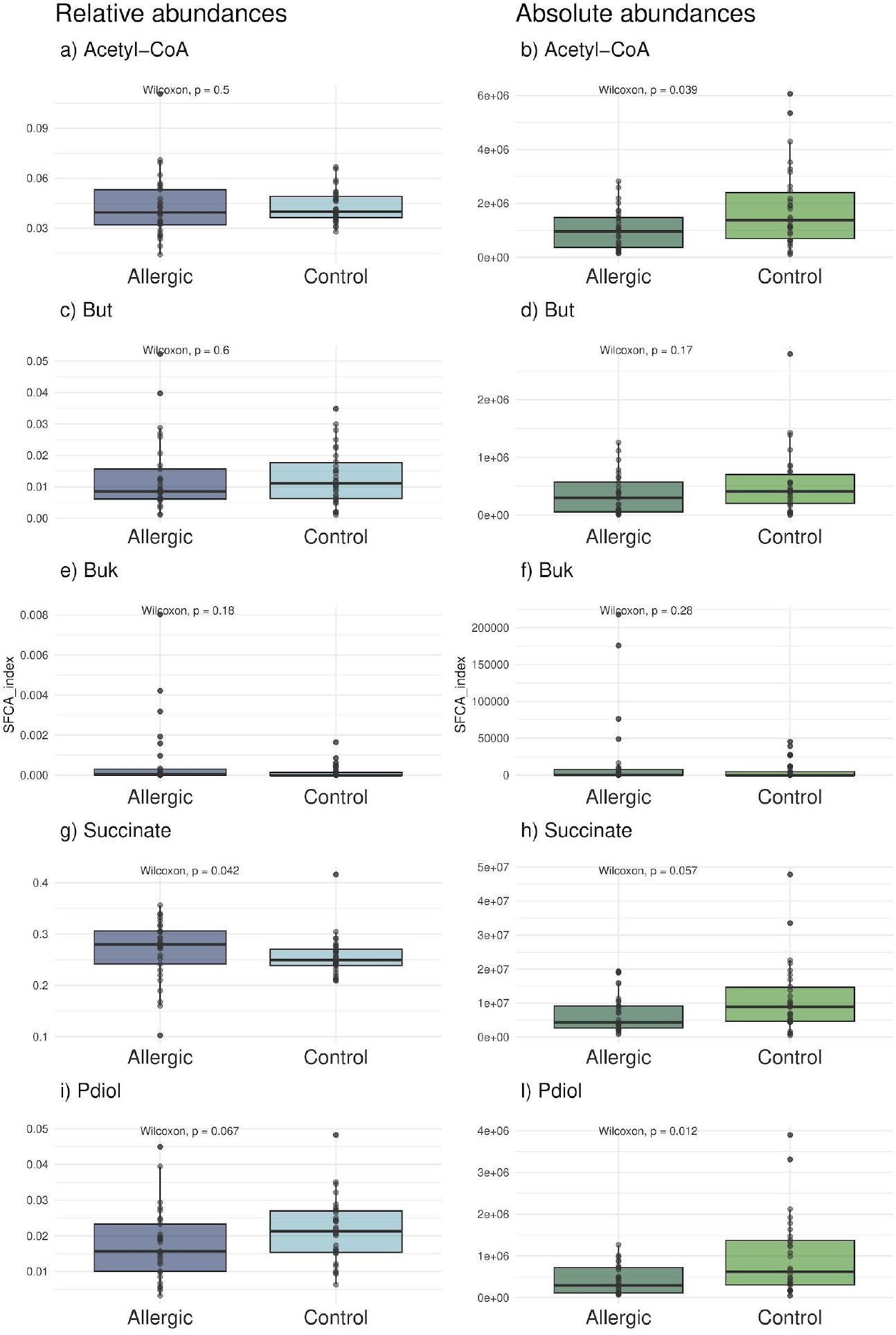
Picrust2 predictions for short chain fatty acids (SCFAs) production in control and allergic children using relative (a, c, e, g and i, on the left) and absolute (b, d, f, h and l on the right) abundances. Acetyl-Coa = total predicted capacity of producing Acetyl Coenzyme A (butyrate pathway). But = total predicted abundance of taxa carrying the *but* gene (butyrate pathway). Buk = total predicted abundance of taxa carrying the *buk* gene (butyrate pathway). Succinate = total predicted succinate production capacity (propionate pathway). Pdiol = total predicted abundance of genes involved in propanediol utilization to produce propionate. In the boxplots for relative abundances (a, c, e, g and i), allergies are reported in dark blue and control individuals in light blue. Boxplots for absolute abundances (b, d, f, h and l), the allergic group is reported in dark green and control individuals in light green. Abundances for allergic and control children were compared using the Wilcoxon–Mann–Whitney test, with a significance threshold (p-value) set to 0.05.

Concerning butyrate, relative and absolute abundances yielded contrasting predictions for the overall Acetyl-CoA production capacity, a key component of the butyrate synthesis pathway. Based on absolute counts, food-allergic children showed a significantly lower predicted Acetyl-CoA production capacity than controls, whereas this difference was not evident when using relative abundances. In contrast, both relative and absolute abundances consistently indicated similar predicted abundances of taxa carrying the *but* and *buk* genes, which encode the terminal enzymes of butyrate biosynthesis.

Discordant predictions between relative and absolute abundance datasets were also observed for tot_pdiol, which reflects the genomic potential for propanediol-based propionate production. Indeed, a significantly lower predicted abundance of propanediol-utilization genes in allergic children emerged only when absolute bacterial counts were considered. Lastly, for succinate, a precursor in propionate biosynthesis, both datasets showed significant group differences, but with reversed direction. Relative abundances suggested a greater succinate production potential in allergic children, whereas absolute abundances supported the opposite conclusion.

## DISCUSSION

The oral microbiota, a key component of the oral ecosystem, is emerging as a crucial factor associated with the pathogenesis and progression of food allergies. Evidence is accumulating on the alterations of oral bacterial community structure between patients and healthy counterparts ^[24,25]^.

Unfortunately, for technical limitations most of the investigations on microbiota composition are based on relative taxa frequencies, producing biased results and limiting our understanding of the complex interaction between host and the microbial community ^[14]^.

In a previous work we found that the oral microbiota significantly differ in children with food allergy, in comparison to controls, on the basis of relative taxa frequencies determined by standard 16S V3-V4 rRNA amplicon sequencing ^[11]^. In order to better explore the role of oral microbiota in food allergy, we decided to re-analyze the DNA samples from our previous study^[11]^ to determine the absolute abundances of taxa.

We found that allergic children harbor a significantly lower bacterial load in their saliva, in comparison to controls, a pivotal feature impossible to emerge *via* standard 16S rRNA amplicon sequencing treatment, which only returns relative abundances. This reduction in total bacterial loads may reflect the immunological milieu characteristic of food allergies, including the T helper 2 (Th2)–skewed immune profile and associated disturbances in mucosal homeostasis. For example, alterations in epithelial barrier function, in profiles of salivary antimicrobial peptides, in oral microbial metabolites (SCFAs), and in secretory IgA responses have all been reported in allergic conditions ^[24,26]^. Secretory IgA, in particular, could be implied in the observed reduction of the bacterial abundance. Indeed they are known to play a central role in shaping microbial community structure, regulating both the abundance and diversity of commensal taxa and mediating host–microbe interactions ^[27,28]^. Interestingly, perturbations in IgA responses, reported in allergies and in autoimmune diseases, have been associated with impaired immune tolerance toward commensals at mucosal surfaces and disruption of bacterial network stability ^[29,30]^. Moreover, in a mouse model of food allergy characterized by dysregulated IgA production, a significant decrease in the absolute abundance of a dominant oral genus, *Lactobacillus* spp., has been documented ^[30]^, supporting the possible link between altered IgA function and shifts in oral microbial ecology. In addition to this, the low-grade mucosal inflammation reportedly associated with food allergy may contribute to modifying the nutrient and physicochemical environment of saliva, impacting the ability of the oral mucosa to support or control microbial colonization, and thus the total bacterial load ^[24]^.

Coherently with our results, reduced total loads in bacterial communities of various body districts have already been reported for other immune-related diseases. Specifically, previous data reported a reduced bacterial colonization in the fecal microbiota of patients with Crohn’s disease ^[31]^ and in the salivary microbiota of children with high salivary glucose concentration, at risk for Type 2 Diabetes ^[32]^. Also these works conclude that the estimation of the microbial load of samples is a key point for evaluating microbiota alterations. Moreover both papers raise the possibility that altered overall abundance itself could be a key feature of a disease-associated microbiota configuration. Hence, the possibility of using overall, or taxa specific, abundances for diagnosis and/or outcome predictive purposes ^[32]^. Specifically, Goodson and colleagues ^[32]^ observe that the determination of total salivary microbial loads was able to predict high oral glucose concentration and obesity-related metabolic syndrome (Type 2 diabetes) more accurately than did clinical measures ^[32,33]^.

Considering the observed variation in absolute bacterial loads between allergic and control groups, we decided to combine total bacterial load with taxa relative abundances to obtain absolute abundances of all taxa identified in our previous work (D’Auria et al. 2023). For this analysis, we followed a pipeline already suggested by Bruijning and colleagues (2023) ^[14]^, which also highlighted how the use of relative taxa abundances can bias microbiota analyses. The obtained absolute abundances were then used to re-evaluate β-diversity indices, taxonomic pictures and SCFA production capacity, then compared with those from the relative abundances.

Regarding β-diversity, the PERMANOVA on the PCoA continued to show a significant separation between allergic and control children when using the absolute abundances.

For what concerns the taxonomic profiles, the use of absolute abundances has partially reshaped our previous findings in D’Auria et al. (2023) ^[11]^. About one fourth of the taxa displayed significant incoherent relative or absolute abundance between allergic and control children. The relative abundances of *Patescibacteria* and unclassified lineages appear artefactually increased in allergic children because their absolute abundances remain essentially unchanged, while the overall bacterial load is markedly reduced in this group. For the same reason, the reduction of *Actinobacteriota, Bacteroidota* and *Spirochetota* in allergic children, apparent from absolute abundances, is obscured when relying solely on relative-frequency comparisons. The case of *Firmicutes* appears unique: their significant reduction in allergic children is detected by both relative and absolute quantifications. This likely reflects a particularly pronounced decrease, greater than that observed for the other bacterial lineages, such that it remains detectable even in the context of the overall reduction in total bacterial load in the allergic group.

The differential contribution of individual phyla to the overall reduction in salivary bacterial loads raises several noteworthy points. Most of the major bacterial phyla inhabiting the oral cavity are affected by this decrease, which reaches its maximum for *Firmicutes*. This finding is consistent with current knowledge on the immunological landscape of allergic diseases and the role played by *Firmicutes* therein ^[24,26]^. Analyses at finer taxonomic levels confirmed a marked reduction in key *Firmicutes* taxa with known anti-inflammatory properties, including SCFA-producing families such as *Lachnospiraceae, Ruminococcaceae*, as well as taxa correlated with local Th1 cytokine production in the oral cavity, such as *Streptococcus* ^[30]^. Notably, depletion of *Firmicutes* has been reported as the principal taxonomic shift characterizing oral dysbiosis resulting from perturbations in secretory IgA responses ^[28]^, which have also been described in allergic conditions (^[30]^ and above). *Patescibacteria*, along with sequences unresolved by the 16S rRNA V3-V4 amplicon sequencing represent the portion of the oral microbiota of allergic children that remains stable in absolute quantity, even in the context of the marked reduction of the oral bacterial load. For this reason, these taxa appear artefactually increased in allergic children when considering the relative frequencies. The fact that these bacterial lineages are not affected by the perturbations in the oral environment that lead to the significant decrease observed for several major bacterial phyla is intriguing. This could be related to particular ecological niches occupied by these bacteria, to specific features of their lifestyle or metabolism allowing them to adapt to the new conditions, or to particular interactions they may have with the immune system ^[26]^.

With respect to *Patescibacteria*, our observations are consistent with reports describing increased relative abundances of *“Ca*. Saccharimonadia” (the most extensively studied *Patescibacteria* lineage) in dysbiotic microbiomes and inflammatory environments. Such increases have been documented both in oral pathologies (e.g., periodontitis and gingivitis) and in systemic inflammatory conditions, including Inflammatory Bowel Disease (IBD) and vaginosis ^[12]^. “*Ca*. Saccharimonadia” have indeed been described as “inflammophilic bacteria” capable of thriving in inflammation-associated, nutrient-rich niches ^[13]^. Notably, “*Ca*. Saccharimonadia” also exert important immunomodulatory functions, including the regulation of cytokine responses and the immunoactivation of oral epithelial cells ^[34,35]^. Furthermore, recent evidence indicates that “*Ca*. Saccharimonadia” can be endocytosed by oral epithelial cells, and survive long-term, a mechanism that may at least partly account for their unchanged absolute abundances in our work ^[35]^. Taxonomic characterization of the unclassified bacterial sequences, representing the other fraction unaffected by the overall bacterial decrease, revealed that they comprise a heterogeneous set of low-abundance lineages spanning all major bacterial phyla known to inhabit the oral cavity ^[26]^. Their heterogeneity and broad representation across bacterial diversity may suggest that factors other than simple taxonomic affiliation, such as niche specialization or interactions with the host immune system, contribute to their persistence.

Finally, our results showed that the biases introduced by the comparison of relative abundances also extend to the prediction of community functional features, such as the inferred composition in SCFA-producing bacteria. Only analyses based on absolute abundance data revealed that allergic children harbored an oral microbiota with a reduced abundance of taxa predicted to produce succinate, as well as lower abundances of taxa involved in propanediol (Pdiol, a propionate precursor) production and of taxa generating butyrate via the acetyl-CoA pathway. Notably, the distortion introduced by relative-abundance-based inference was sufficiently pronounced that succinate production potential appeared significantly increased in the allergic group when assessed using relative data alone. Importantly, predictions derived from absolute abundance measurements are consistent with the taxonomic patterns described above, particularly the depletion of *Firmicutes*, and with previous experimental measurements of salivary SCFA levels in food-allergic children compared with controls ^[10]^.

One of the main limits of this study is the approximate, and potentially biased, taxonomic resolution of 16S rRNA V3-V4 amplicon sequencing. In the future, this limitation could be tackled using shotgun metagenomics for bacterial taxa assignment.

## CONCLUSIONS

In this work we investigated the oral microbiota of allergic children, also taking in account the inherent problems linked to the statistical analysis of interdependent data, as relative abundance data in amplicon-based studies ^[10]^. Recent studies showed that the using of relative abundances of microbes can lead to false conclusions, thus highlighting the importance of absolute bacterial taxa quantification to alleviate this bias ^[19]^. Nevertheless, most 16S rRNA amplicon–based microbiome studies continue to rely exclusively on relative taxon abundances, and few studies have compared inferences derived from absolute versus relative abundance data ^[18,19,36]^. The present work has confirmed that substantial variation of the bacterial load across samples hinders efforts to associate microbiota features with the phenotype under investigation by forcing spurious statistical trends. Moreover, this study uncovers a key biological characteristic of children with food allergy that had remained undetected in the previous, standard, analyses ^[11]^. Specifically, children with food allergy exhibit significantly reduced total oral bacterial loads. This observation is consistent with previous reports in Crohn’s disease ^[31]^ and Type 2 Diabetes ^[32]^, and it adds to the list a disease that is experiencing a dramatic increase in prevalence. Moreover, these findings strengthen the notion that alterations in overall microbiota abundance may themselves represent a defining feature of disease-associated ecosystem configurations, with potential predictive and diagnostic relevance ^[31–33]^.

## DECLARATIONS

## Acknowledgments

We want to thank the Romeo ed Enrica Invernizzi Foundation for support. The authors wish also to thank the support of the APC Central Fund of the University of Milan.

## Authors’ contributions

SP: planned and designed the study, performed molecular biology experiments and drafted and revised the manuscript; LS: performed bioinformatic analyses; CB: performed molecular biology experiments; MA: contributed to design the study; ECD: contributed to design the study and drafted the manuscript; GZ: contributed to design the study and drafted the manuscript; FC: planned and designed the study, performed bioinformatic/statistical analyses and drafted and revised the manuscript.

All authors approved the final version of the manuscript.

## Availability of data and materials

All data produced in this work are available in the supplementary material.

## Ethical approval and consent to participate

This study analyzed the collection of human salivary DNA samples already characterized in a previous work, D’Auria et al. 2023. The research was conducted according to the declaration of Helsinki, and all methods followed the relevant guidelines and regulations. The Ethics Committee of ASST-Fatebenefratelli-Sacco approved the study (Ref. n. 2021/ST/041). Privacy rights of subjects were carefully observed and authors did not have access to information that could identify individual participants. During the recruitment of subjects, parents signed a written informed consent to allow their child to participate in the study. Samples and data were first accessed for research purposes on 01 February 2022.

## Consent for publication

Not applicable.

